# SAMP1/TMEM201 protects alveolar type 2 cells from DNA damage-associated senescence and restrains pulmonary fibrosis

**DOI:** 10.64898/2026.11.06.764769

**Authors:** Leonid A Serebryannyy, David A Ball

## Abstract

Idiopathic pulmonary fibrosis (IPF) is a progressive interstitial lung disease in which repetitive injury to the alveolar epithelium and senescence of alveolar type 2 (AT2) cells drive persistent myofibroblast activation and extracellular matrix deposition. Inner nuclear membrane (INM) proteins organise nuclear architecture and participate in the DNA damage response, but whether they influence epithelial senescence and fibrogenesis is unknown. Here we identify SAMP1/TMEM201, an INM protein that binds lamin A/C, emerin and LINC-complex components, as a compartment-selective protector of AT2 cells. In the bleomycin model of pulmonary fibrosis, SAMP1 was reduced in AT2 cells but unchanged in lung fibroblasts. AT2 cell–specific deletion of Samp1 (SftpcCre ERT2; Samp1-flox/flox) aggravated fibrosis at day 21, with higher Ashcroft scores and increased pulmonary Tnfa and Acta2 expression, whereas intratracheal delivery of AAV6-CAG-Samp1 before bleomycin attenuated both histological fibrosis and these transcriptional responses. In primary AT2 cells analysed immediately after isolation without in vitro stimulation, SAMP1 loss was accompanied by reduced 53BP1 protein, and upregulation of the senescence-associated and profibrotic markers p16 and PAI-1. These findings define a SAMP1–AT2 senescence axis that constrains pulmonary fibrogenesis and suggest that preserving INM-dependent genome surveillance in the alveolar epithelium may offer an epithelium-directed therapeutic strategy for IPF. Because DNA damage signaling was assessed only under baseline conditions, the data establish an association with altered DNA damage signaling rather than a definitive defect in lesion repair.

## Introduction

Idiopathic pulmonary fibrosis (IPF) is a chronic, progressive and ultimately fatal interstitial lung disease characterized by excessive deposition of extracellular matrix (ECM), destruction of alveolar architecture and irreversible loss of lung function, with a median survival of only 3–5 years after diagnosis [1,2,40]. Although the approved antifibrotic agents pirfenidone and nintedanib — and, more recently, the PDE4B inhibitor nerandomilast — slow the rate of functional decline, none reverses established fibrosis, and lung transplantation remains the only definitive intervention [2,3,4,41]. A substantial body of evidence indicates that IPF is initiated by repetitive microinjury to the alveolar epithelium. When epithelial repair fails, persistent activation of resident fibroblasts and their differentiation into α-smooth muscle actin (α-SMA)-positive myofibroblasts drives progressive scarring [5,8]. Bleomycin (BLM), a glycopeptide that generates DNA strand breaks, remains the most widely used experimental inducer of pulmonary fibrosis precisely because it reproduces this epithelial injury-driven sequence in mice [6].

Alveolar type 2 (AT2) cells are the facultative progenitors of the alveolar epithelium. They synthesize and secrete pulmonary surfactant and serve as stem cells that replenish both AT2 and type 1 (AT1) cells after injury, thereby occupying a central position in alveolar homeostasis and repair [7,40]. Increasingly, dysfunction of AT2 cells — rather than of fibroblasts or infiltrating immune cells — is regarded as the initiating event in lung fibrogenesis [5,8]. AT2 cells from IPF lungs, and from BLM-injured mouse lungs, show shortened telomeres, persistent DNA damage signalling and a senescence-like phenotype characterized by induction of the cyclin-dependent kinase inhibitors p16^INK4a^ and p21 [9,10,42]. Critically, AT2 cell senescence is an active driver rather than a passive correlate of fibrosis. Induction of p53-dependent senescence selectively in adult mouse AT2 cells, through deletion of Sin3a, is sufficient to produce spontaneous and progressive pulmonary fibrosis, and the phenotype is abrogated when senescence is blocked or senescent cells are cleared [8,42]. Likewise, AT2 cell–specific deletion of the shelterin component TRF1 causes telomere dysfunction, increases γH2AX DNA damage foci and leads to age-dependent lung remodeling and fibrosis [11,43]. Senescent AT2 cells elaborate a senescence-associated secretory phenotype (SASP) that includes transforming growth factor-β (TGF-β), interleukin-6 (IL-6) and plasminogen activator inhibitor-1 (PAI-1). These factors act in an autocrine and paracrine manner to promote myofibroblast differentiation and ECM accumulation [10,12,13,42]. Among them, PAI-1 is of particular interest: it is not merely a marker but a functional mediator of senescence, inducing p16 and being required for TGF-β1–induced AT2 cell senescence and SASP secretion, whereas PAI-1 deletion or pharmacological inhibition attenuates BLM-induced fibrosis [12,44,45].

Persistent DNA damage signalling is the proximal trigger for such senescence programmes, and the efficiency with which DNA double-strand breaks (DSBs) are sensed and repaired determines whether an injured epithelial cell re-enters the cell cycle or commits to stable arrest [9,10,42]. This response is organized in a spatially structured nuclear environment. DSBs are marked by phosphorylation of the histone variant H2AX (γH2AX), which recruits mediator proteins such as 53BP1 that govern repair pathway choice and checkpoint activation [14,15]. Notably, DNA repair is tightly coupled to the nuclear envelope: chromatin domains anchored at the inner nuclear membrane (INM) display distinct repair kinetics, damaged loci relocate to nuclear pore complexes, and LINC (linker of nucleoskeleton and cytoskeleton) complex components connect the nucleus to cytoskeletal motors to mobilize broken chromosomes [16,17,46]. Accordingly, perturbation of INM proteins — including SUN-domain proteins and A-type lamins — delays DSB repair, causes accumulation of DNA damage and produces premature ageing phenotypes, and mutations affecting INM constituents give rise to laminopathies and progeroid syndromes marked by persistent γH2AX foci and cellular senescence [18,46]. These observations raise the possibility that INM proteins constitute a previously unrecognized class of regulators of epithelial senescence and fibrosis.

SAMP1, also known as TMEM201, is an evolutionarily conserved multi-pass transmembrane protein of the INM. Its nucleoplasmically exposed N-terminal domain contains CxxC zinc-finger motifs that mediate INM targeting and that bind lamin A/C, emerin and the LINC complex components SUN1 and SUN2, as well as the small GTPase Ran [19,20,21]. Through these interactions SAMP1 contributes to nuclear envelope organization, nuclear migration during cell polarization, mitotic spindle assembly and peripheral heterochromatin anchoring [22,23,50]. Loss of SAMP1 impairs endothelial cell migration and angiogenesis in vitro and in vivo [24], and in malignant cells SAMP1 has been reported to positively regulate TGF-β/SMAD2/3 signalling and to promote epithelial–mesenchymal transition, invasion and chemoresistance [25,26]. Despite these advances — including a recent preprint extending SAMP1 biology to hepatic metabolic regulation [49] — the role of SAMP1 in the lung is entirely unknown — in particular, whether this INM protein participates in the DNA damage response and senescence of AT2 cells, or in the pathogenesis of pulmonary fibrosis.

In the present study we addressed this question. We show that SAMP1 expression is selectively reduced in AT2 cells, but not in lung fibroblasts, of mice with BLM-induced pulmonary fibrosis (Figure 1). AT2 cell–specific deletion of Samp1 exacerbates BLM-induced lung fibrosis, with increased histological fibrosis scores and elevated pulmonary expression of Tnfα and Acta2 (Figure 2), whereas AAV-mediated restoration of SAMP1 expression in the lung attenuates these responses (Figure 3). Mechanistically, primary AT2 cells lacking SAMP1 display a compromised DNA damage response, as evidenced by reduced 53BP1 protein levels, together with upregulation of the senescence and profibrotic markers p16 and PAI-1 (Figure 4). Collectively, these findings identify SAMP1 as a protector of AT2 cells against DNA damage– induced senescence and establish the SAMP1–AT2 senescence axis as a contributor to pulmonary fibrogenesis, with implications for therapeutic strategies aimed at preserving the alveolar epithelium.

**Figure 1.**
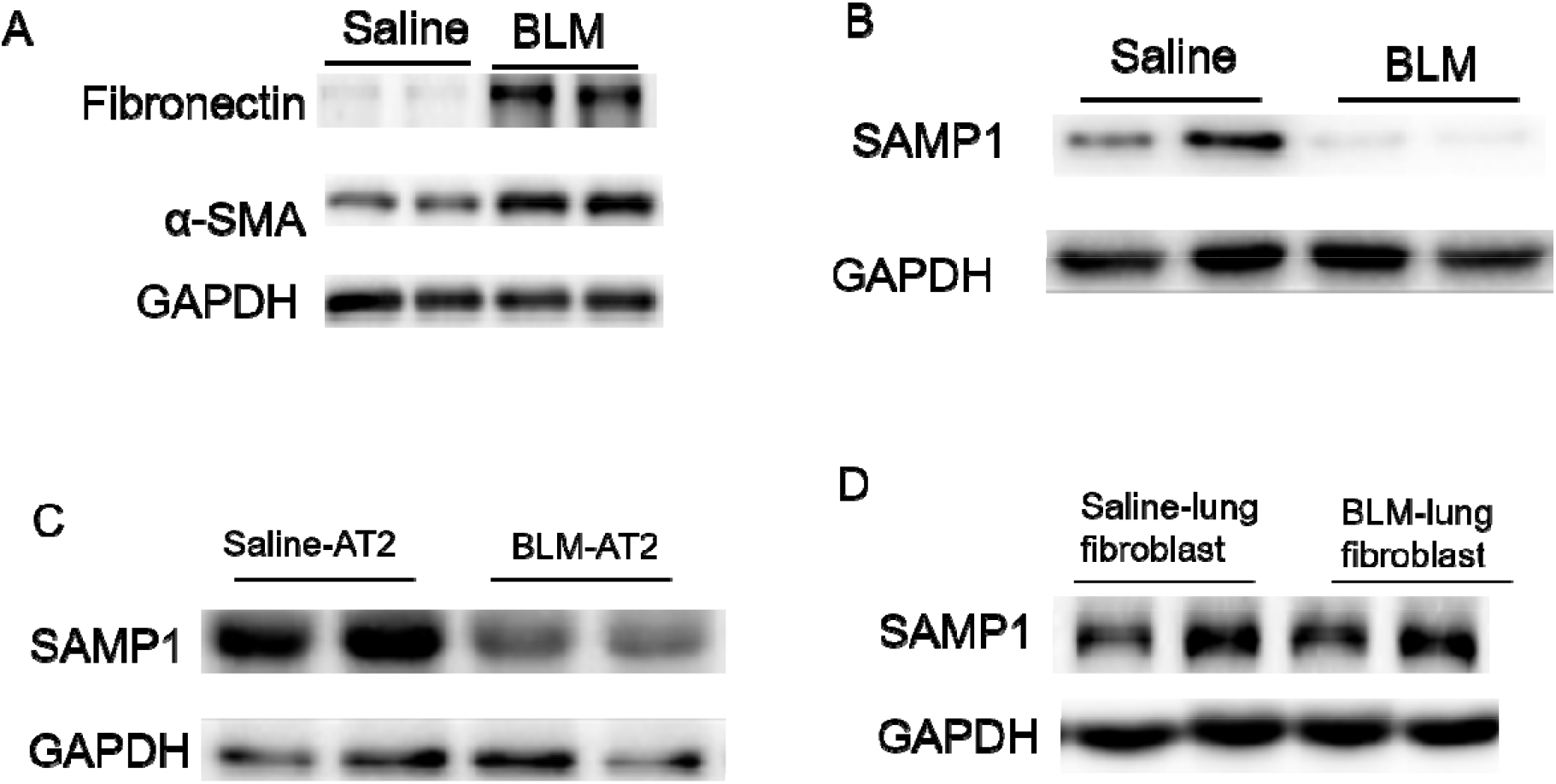
SAMP1/TMEM201 is downregulated in alveolar type II epithelial cells during bleomycin-induced pulmonary fibrosis. (A) Representative Western blots of the fibrotic markers fibronectin and α-SMA in lung tissues from saline- and bleomycin (BLM)-treated mice, confirming successful establishment of the fibrosis model (n = 2 per group shown). GAPDH served as loading control. (B) Western blot of SAMP1 in whole-lung homogenates from saline- and BLM-treated mice. SAMP1 expression was markedly reduced in BLM-challenged lungs. GAPDH served as loading control. (C) Western blot of SAMP1 in primary alveolar type 2 (AT2) epithelial cells isolated from saline- and BLM-treated mice. SAMP1 was significantly downregulated in BLM-injured AT2 cells. GAPDH served as loading control. (D) Western blot of SAMP1 in primary lung fibroblasts isolated from saline- and BLM-treated mice. SAMP1 expression was comparable between groups, indicating that SAMP1 downregulation is enriched in AT2 cells rather than fibroblasts. GAPDH served as loading control.

**Figure 2.**
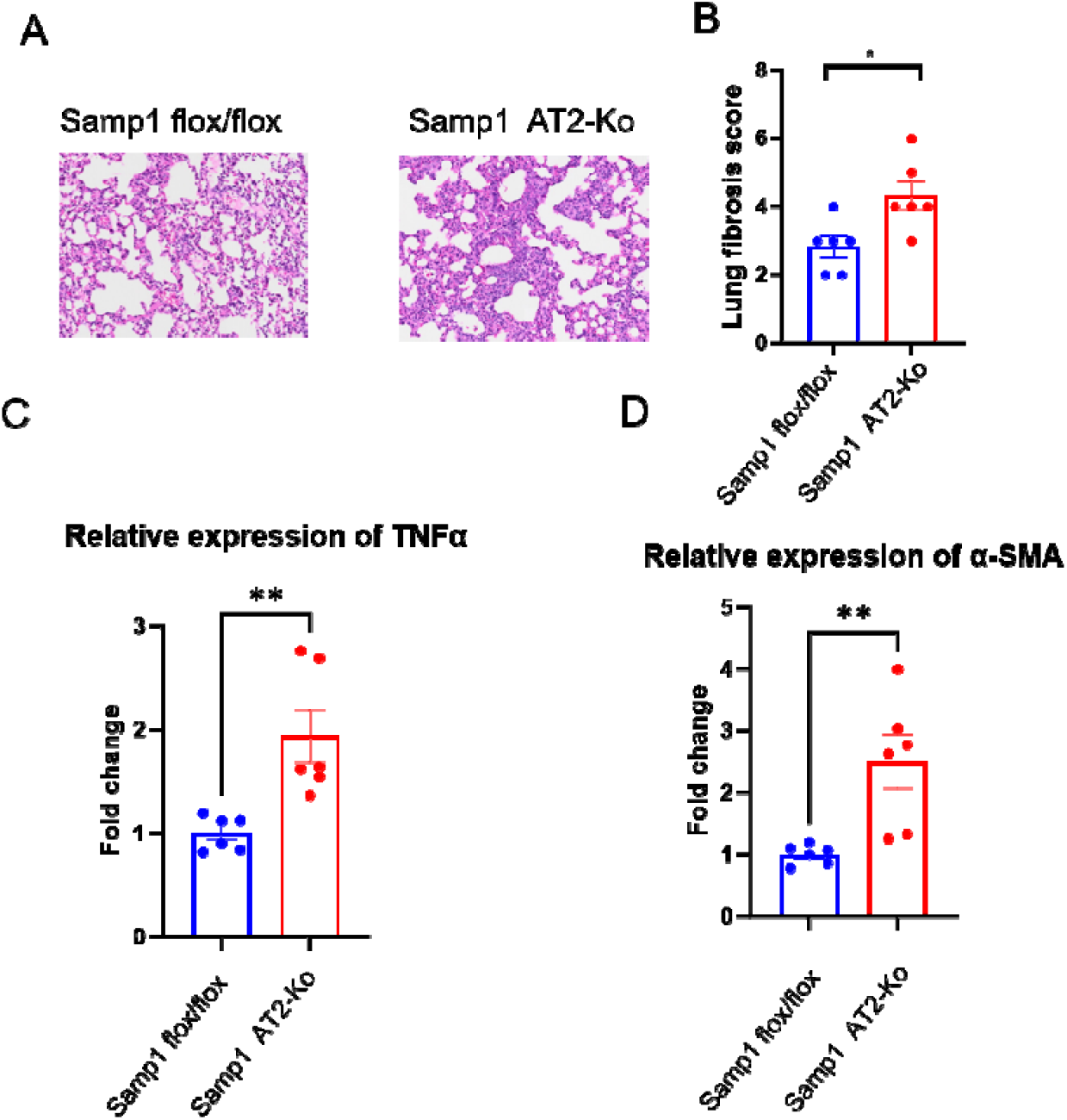
AT2-specific deletion of Samp1 exacerbates bleomycin-induced pulmonary fibrosis. (A) Representative hematoxylin and eosin (H&E) staining of lung sections from Samp1^flox/flox^ and Samp1^AT2-KO^ mice at day 21 after BLM challenge. Scale bar, 100 μm. (B) Quantification of lung fibrosis scores in the two groups (n = 5 per group). Data are presented as mean ± SEM; *P < 0.05 by unpaired two-tailed Student’s t-test. (C) Relative mRNA expression of TNFα in lung tissues, measured by qPCR and normalized to Gapdh. Data are mean ± SEM of n = 5–6 mice per group; **P < 0.01 vs. Samp1^flox/flox^ group. (D) Relative mRNA expression of α-SMA (Acta2) in lung tissues. Data are mean ± SEM of n = 5–6 mice per group; **P < 0.01 vs. Samp1^flox/flox^ group.

**Figure 3.**
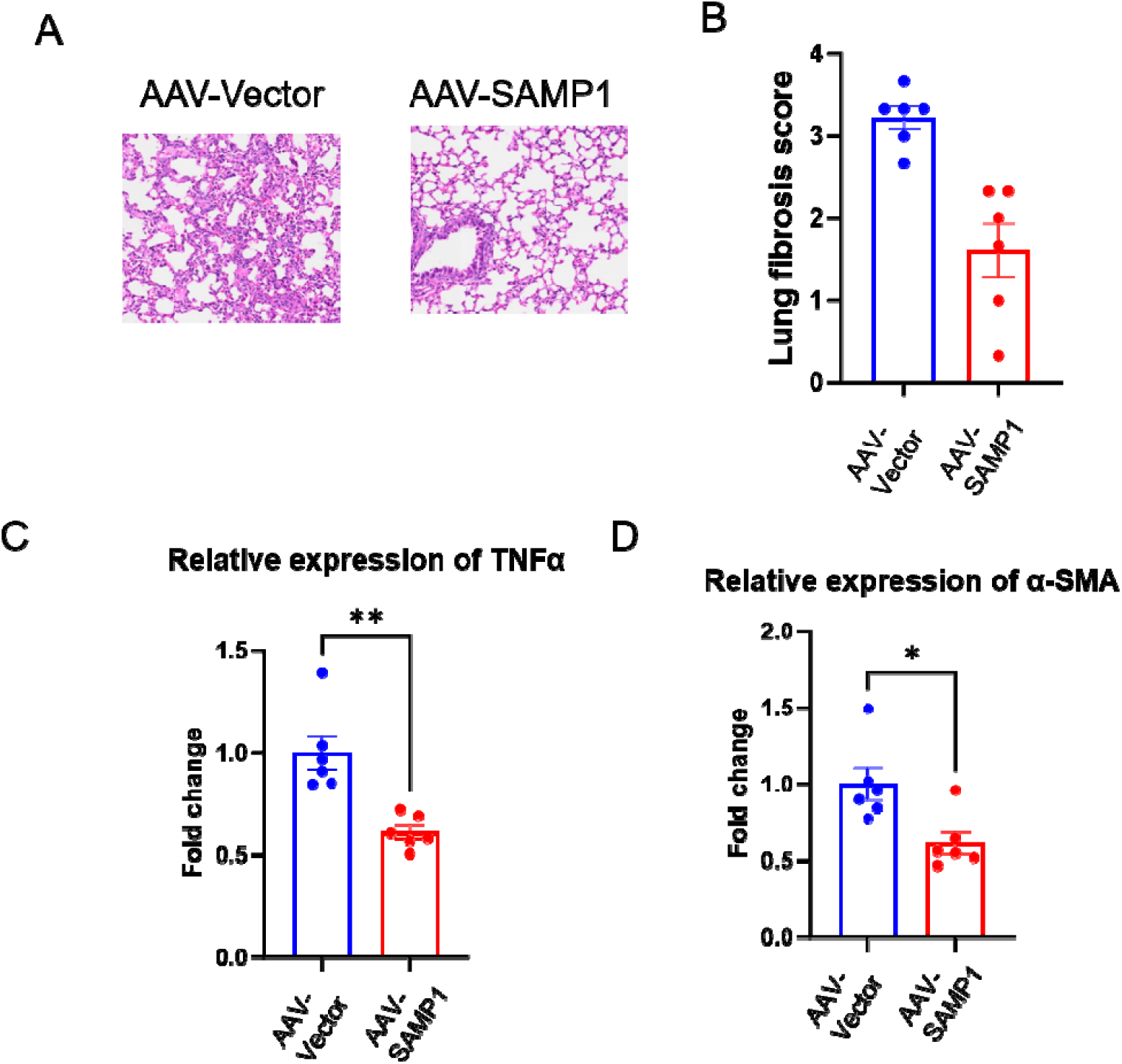
AAV-mediated SAMP1 overexpression attenuates bleomycin-induced pulmonary fibrosis. (A) Representative H&E staining of lung sections from BLM-challenged mice receiving AAV-Vector or AAV-Samp1. Scale bar, 100 μm. (B) Quantification of lung fibrosis scores (n = 5 per group). Data are mean ± SEM; **P < 0.01 vs. AAV-Vector group by unpaired two-tailed Student’s t-test. (C) Relative mRNA expression of TNFα in lung tissues. Data are mean ± SEM of n = 5–6 mice per group; **P < 0.01 vs. AAV-Vector. (D) Relative mRNA expression of α-SMA (Acta2) in lung tissues. Data are mean ± SEM of n = 5–6 mice per group; *P < 0.05 vs. AAV-Vector.

**Figure 4.**
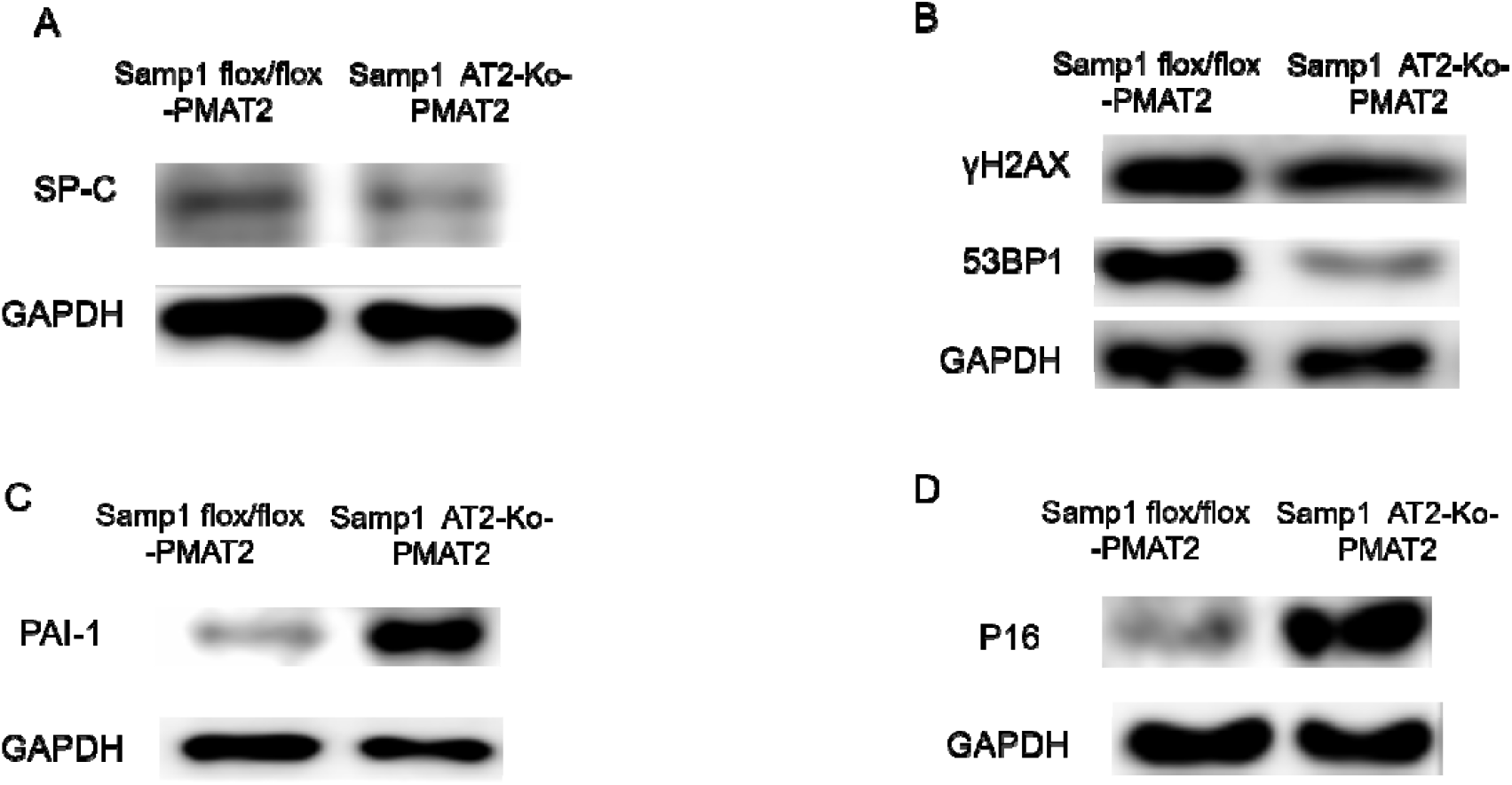
SAMP1-deficient AT2 cells exhibit an impaired DNA damage response and a senescent, profibrotic phenotype. (A) Primary mouse alveolar type 2 (PMAT2) cells were isolated from Samp1^flox/flox^ (WT) and Samp1^AT2-KO^ mice. Representative Western blot of surfactant protein C (SP-C) and GAPDH confirming the AT2 identity of the isolated cells. GAPDH served as loading control. (B) Representative Western blots of γH2AX, 53BP1 and GAPDH in PMAT2 cells of the indicated genotypes. 53BP1 protein levels were decreased in Samp1 AT2O^cells, whereas γH2AX levels were comparable between the two genotypes (C,D) Representative Western blots of PAI-1 (SERPINE1) and p16^INK4a^, with GAPDH as a loading control, in PMAT2 cells of the indicated genotypes. Both PAI-1 and p16^INK4a^ were upregulated in Samp1-deficient AT2 cells. Densitometric quantification of the blots shown in (B)–(D) (right/lower panels). Band intensity was normalized to GAPDH and expressed as fold change relative to WT cells (set as 1.0). Data are presented as mean ± SD from n = 3 independent experiments. *P < 0.05, **P < 0.01, ***P < 0.001; ns, not significant (two-tailed unpaired Student’s t-test). Molecular weights (kDa) are indicated to the right of each blot.

## Materials and Methods

### Animals

Samp1^flox/flox^ mice on a C57BL/6J background were generated by CRISPR/Cas9-mediated insertion of loxP sites flanking the critical coding exons of Tmem201 [targeting vector; floxed exon numbers; vendor; strain no.] and were crossed with Sftpc-Cre-ERT2-mice (The Jackson Laboratory, Bar Harbor, ME; stock no. 028054; RRID: IMSR_JAX:028054) [7,31] to generate Sftpc-Cre^ERT2^; Samp1^flox/flox^ mice, hereafter designated Samp1^AT2-KO^. Littermates carrying the floxed alleles but lacking Cre were used as wild-type (WT) controls. To induce Cre-mediated recombination, mice aged 8–12 weeks received tamoxifen (75 mg/kg body weight; 20 mg/mL in corn oil; Sigma-Aldrich, CAS 10540-29-1) by intraperitoneal injection once daily for 5 consecutive days, followed by a 14-day washout before any further intervention; WT littermates received the same tamoxifen regimen [32]. Genotyping was performed by PCR on tail genomic DNA using Samp1 flox forward/reverse and Cre forward/reverse primer pairs (sequences, product sizes and annealing temperatures are listed in Supplementary Table 1). Both male and female mice were used, and animals were age- and sex-matched across experimental groups. Mice were housed under specific pathogen–free conditions on a 12 h light/12 h dark cycle with free access to food and water. All animal procedures were reviewed by the Animal Welfare Body (Organismo Preposto al Benessere Animale, OPBA) of Sapienza University of Rome (Sapienza Università di Roma) and authorized by the Italian Ministry of Health (authorization no. 721/2022-PR) in accordance with Legislative Decree No. 26/2014 implementing Directive 2010/63/EU; the study was conducted and reported in accordance with the ARRIVE 2.0 guidelines [33].

### Bleomycin-induced pulmonary fibrosis model

Mice were anesthetized with inhaled isoflurane (2–3% for induction and 1.5–2% for maintenance in oxygen) and a single dose of bleomycin sulfate (1.5 U/kg body weight, equivalent to approximately 1.6 mg/kg; Nippon Kayaku, Tokyo, Japan) dissolved in 50 μL of sterile saline was administered by intratracheal instillation using a blunt 22-G catheter. Control mice received an equal volume of sterile saline. Body weight and general condition were monitored daily throughout the experiment. Mice were euthanized on day 21 after instillation, and lungs were harvested for histological, biochemical and molecular analyses [6,34,48].

### AAV-mediated Samp1 overexpression in vivo

Recombinant adeno-associated virus vectors of serotype AAV6 encoding mouse Samp1 (AAV-Samp1), or the corresponding empty vector control (AAV-Vector), were constructed with the transgene under the control of the constitutive CAG promoter and produced by InnovAAVector s.r.l. (Pozzuoli, Naples, Italy) at a titre of 2 × 10^12 vg/mL. Mice were anesthetized as described above and received 1 × 10^11 vg of AAV-Samp1 or AAV-Vector in 50 μL of sterile saline by intratracheal instillation 14 days before bleomycin administration [30,35,36,47]. Transduction efficiency and transgene expression were verified at the time of harvest by Western blotting of whole-lung lysates and by immunofluorescence co-staining of SAMP1 with pro-SP-C on lung sections.

### Histological analysis and fibrosis scoring

The left lung was fixed in 4% paraformaldehyde for 24 h at room temperature, embedded in paraffin and sectioned at 4 μm. Sections were stained with hematoxylin and eosin (H&E) according to standard protocols. Images were acquired on a bright-field microscope [model; camera] using a ×20 objective (scale bar, 100 μm). Fibrosis was scored independently by two investigators blinded to genotype and treatment using the modified Ashcroft scoring system [27,28] over 20 randomly selected non-overlapping fields per mouse, and the mean score per mouse was used for statistical analysis. Where indicated, collagen deposition was further assessed by Masson’s trichrome staining, and total lung collagen was quantified by measuring hydroxyproline content with a commercial hydroxyproline assay kit (Sigma-Aldrich, MAK0008, St. Louis, MO) after acid hydrolysis of lung homogenates in 6 N HCl at 120 °C for 3 h, according to the manufacturer’s instructions [39].

### Isolation of primary mouse alveolar type 2 cells

Primary mouse alveolar type 2 (PMAT2) cells were isolated from Samp1^flox/flox^ and Samp1^AT2-KO^ mice 2 weeks after tamoxifen administration, or from saline- and BLM-treated mice on day 21, essentially as described [29,37]. Briefly, mice were euthanized and the pulmonary circulation was flushed with ice-cold PBS via the right ventricle. Lungs were inflated through the trachea with dispase II (2 mg/mL, approximately 50 U/mL; Sigma-Aldrich) in DMEM/F12 containing 0.01% DNase I, minced, and digested at 37 °C for 45 min with gentle agitation. The digest was sequentially filtered through 100-, 40- and 20-μm cell strainers and centrifuged at 300 × g for 5–10 min at 4 °C. Where necessary, erythrocytes were removed by brief incubation in ACK lysis buffer. Single-cell suspensions were depleted of leukocytes and endothelial cells using biotinylated anti-CD45, anti-CD31, anti-CD16/32 and anti-Ter119 antibodies together with streptavidin-conjugated magnetic beads, after which EpCAM-positive epithelial cells were positively selected with anti-EpCAM (CD326) microbeads. AT2 identity and purity were confirmed by pro-SP-C immunofluorescence on cytospins and routinely exceeded 95% (Figure 4A). Cells were lysed immediately for protein analysis without in vitro stimulation.

### Isolation and culture of primary mouse lung fibroblasts

Primary lung fibroblasts were isolated from saline- and BLM-treated mice on day 21 after instillation. Lungs were minced and digested in 1 mg/mL collagenase type I (Sigma-Aldrich, C0130) in Hanks’ balanced salt solution at 37 °C for 30 min with gentle agitation. The digest was filtered, washed, and plated in DMEM supplemented with 10% FBS and 1% penicillin–streptomycin. Non-adherent cells were removed after 24 h and adherent fibroblasts were expanded; cells at passage 2–3 were used for all experiments to avoid confounding effects of culture-induced replicative senescence. Purity was verified by positive staining for vimentin and PDGFRα and absence of EpCAM and CD45 expression.

### Western blot analysis

Lung tissues and cells were lysed in RIPA buffer supplemented with protease and phosphatase inhibitor cocktails, and protein concentration was determined by the BCA assay. Equal amounts of protein (30 μg per lane) were separated by SDS-PAGE on 10% gels and transferred to PVDF membranes. Membranes were blocked with 5% non-fat milk or bovine serum albumin in Tris-buffered saline containing 0.1% Tween-20 for 1 h at room temperature, incubated with primary antibodies overnight at 4 °C, and then with horseradish peroxidase–conjugated secondary antibodies for 1 h at room temperature. Immunoreactive bands were visualized by enhanced chemiluminescence (ECL substrate [supplier; catalogue no.]) and quantified by densitometry using ImageJ (National Institutes of Health). Target protein levels were normalized to GAPDH and expressed as fold change relative to the control group. For γH2AX, normalization was likewise performed to GAPDH; normalization to total H2AX or histone H3 was not performed (see Limitations).

### Quantitative real-time PCR

Total RNA was extracted from lung tissue or cultured cells using TRIzol reagent (Thermo Fisher Scientific, 15596026CN), and 1 μg of RNA was reverse-transcribed with a commercial cDNA synthesis kit (PrimeScript RT Master Mix, Takara Bio. Quantitative PCR was performed on an Applied Biosystems QuantStudio 3 Real-Time PCR System (Thermo Fisher Scientific) using SYBR Green chemistry under the following cycling conditions: 95 °C for 10 min; 40 cycles of 95 °C for 15 s and 60 °C for 60 s. Melting-curve analysis confirmed amplification specificity. Relative mRNA expression was calculated by the 2^−ΔΔCt^ method with Gapdh as the reference gene [38]. Primer sequences are listed in Supplementary Table 1.

### Antibodies and reagents

The following primary antibodies were used: anti-SAMP1/TMEM201, anti-fibronectin, anti-α-SMA, anti-53BP1, anti-γH2AX, anti-PAI-1 (SERPINE1), anti-p16, anti-pro-SP-C and anti-GAPDH (suppliers, catalogue numbers, RRIDs, host species and working dilutions are listed in Supplementary Table 2). Horseradish peroxidase–conjugated goat anti-rabbit and goat anti-mouse IgG secondary antibodies were purchased from [vendor]. Other reagents were obtained from [vendor] unless stated otherwise.

### Statistical analysis

All quantitative data are presented as mean ± SEM for in vivo experiments and mean ± SD for in vitro experiments, derived from at least three independent experiments. Normality was assessed with the Shapiro–Wilk test and homogeneity of variance with Levene’s test. Comparisons between two groups were performed using an unpaired two-tailed Student’s t-test; comparisons among three or more groups were performed using one-way ANOVA followed by Tukey’s post hoc test, or two-way ANOVA followed by Šídák’s multiple-comparisons test where genotype and treatment were analysed as two factors. Statistical analyses were conducted with GraphPad Prism version 10.1 (GraphPad Software, San Diego, CA, USA). A two-sided P value < 0.05 was considered statistically significant.

## Results

### SAMP1 is downregulated in alveolar type 2 cells in the bleomycin model of pulmonary fibrosis

To establish the fibrosis model, mice received a single intratracheal instillation of bleomycin (BLM) and lungs were harvested on day 21. Compared with saline-treated controls, BLM-challenged lungs showed markedly increased protein expression of the fibrotic markers fibronectin and α-smooth muscle actin (α-SMA), confirming successful induction of pulmonary fibrosis (Figure 1A). We next examined SAMP1 expression in this model. In whole lung homogenates, SAMP1 protein levels were markedly reduced in BLM-treated mice relative to saline-treated controls (Figure 1B). Because the lung is composed of multiple cell types and a change in whole-lung lysates may simply reflect altered cellular composition, we isolated the principal cell populations of interest. In primary mouse alveolar type 2 (PMAT2) cells freshly isolated from BLM-treated mice, SAMP1 was significantly downregulated compared with cells from saline-treated mice (Figure 1C). In contrast, SAMP1 expression in primary lung fibroblasts isolated from the same animals was comparable between the two groups (Figure 1D). These results indicate that the reduction of SAMP1 in the fibrotic lung is not a uniform response of all resident cells but is enriched in, and may be relatively selective for, the AT2 epithelial compartment.

### AT2-specific deletion of Samp1 exacerbates bleomycin-induced pulmonary fibrosis

To determine whether the loss of SAMP1 in AT2 cells is causally linked to fibrogenesis, we generated AT2 cell–specific Samp1 knockout mice by crossing Samp1^flox/flox^ animals with Sftpc-Cre^ERT2^ drivers (Samp1^AT2-KO^) and challenged them with BLM. On day 21 after instillation, hematoxylin and eosin (H&E) staining revealed more extensive alveolar wall thickening, architectural distortion and inflammatory infiltration in Samp1^AT2-KO^ lungs than in Samp1^flox/flox^ littermate controls (Figure 2A). Blinded quantification using a modified Ashcroft scoring system showed a significantly higher fibrosis score in Samp1^AT2-KO^ mice (n = 5 per group; Figure 2B). Consistent with the histological findings, quantitative PCR of lung tissue demonstrated significantly elevated mRNA expression of the proinflammatory cytokine TNFα (Figure 2C) and of the myofibroblast marker α-SMA (Acta2) (Figure 2D) in Samp1^AT2-KO^ mice compared with controls (n = 5–6 per group). Thus, deletion of Samp1 in AT2 cells is sufficient to worsen BLM-induced lung fibrosis and to amplify both the inflammatory and the myofibroblast-associated transcriptional response.

### AAV-mediated SAMP1 overexpression attenuates bleomycin-induced pulmonary fibrosis

Having observed that loss of SAMP1 aggravates fibrosis, we asked whether restoring SAMP1 expression could produce the opposite effect. Mice received AAV-Samp1 or AAV-Vector (1 × 10^11 vg in 50 μL) by intratracheal instillation 14 days before BLM challenge. Overexpression of SAMP1 was confirmed at the time of harvest by Western blotting of whole-lung lysates and by SAMP1/pro-SP-C co-immunofluorescence (Figure 3A and Supplementary Figure 1). H&E staining showed that AAV-mediated SAMP1 delivery visibly preserved alveolar architecture and reduced the extent of fibrotic remodeling relative to the AAV-Vector group (Figure 3A). Quantification confirmed a significant reduction in fibrosis score (n = 5 per group; Figure 3B), accompanied by significantly lower pulmonary mRNA expression of TNFα (Figure 3C) and Acta2 (Figure 3D) (n = 5–6 per group). Together with the genetic loss-of-function data, these gain-of-function results indicate that SAMP1 acts as a protective factor against BLM-induced pulmonary fibrosis, and that augmenting SAMP1 expression in the lung is sufficient to blunt the fibrotic response.

### SAMP1-deficient AT2 cells exhibit an impaired DNA damage response and a senescent, profibrotic phenotype

To explore the cellular basis of the pro-fibrotic effect of SAMP1 loss, we isolated PMAT2 cells from Samp1 flox/flox and Samp1AT2-KO mice and analysed them directly, thereby avoiding confounding contributions from fibroblasts, immune cells or the extracellular environment. Expression of surfactant protein C (SP-C) was readily detected in cells of both genotypes, confirming the AT2 identity and purity of the preparations (Figure 4A). Because SAMP1 is an inner nuclear membrane protein and the nuclear envelope contributes to the organization of DNA repair, we assessed two core components of the DNA double-strand break response. The level of 53BP1, a mediator that accumulates at sites of DNA damage and governs repair pathway choice and checkpoint activation, was decreased in Samp1AT2-KO PMAT2 cells, whereas γH2AX levels were comparable between the two genotypes (Figure 4B). Notably, AT2 cells lacking SAMP1 also displayed a pronounced increase in PAI-1 (SERPINE1) (Figure 4C), a canonical TGF-β target and an established marker and mediator of cellular senescence, and in p16^INK4a^ (Figure 4D), a cyclin-dependent kinase inhibitor that enforces stable cell-cycle arrest and is one of the most widely used markers of senescence. Collectively, these findings indicate that SAMP1 deficiency in AT2 cells is associated with an attenuated DNA damage response machinery together with the induction of a senescent, SASP-associated phenotype, providing a plausible cellular mechanism through which loss of SAMP1 in the alveolar epithelium promotes pulmonary fibrosis.

## Discussion

In this study we identify SAMP1/TMEM201 as a previously unrecognized protective factor in the alveolar epithelium during pulmonary fibrosis. Four lines of evidence support this conclusion. First, SAMP1 is downregulated in AT2 cells, but not in lung fibroblasts, of mice with BLM-induced fibrosis, indicating that the change is compartment-selective rather than a global consequence of tissue injury. Second, AT2 cell–specific deletion of Samp1 exacerbates BLM-induced fibrosis, with higher fibrosis scores and increased pulmonary expression of Tnfα and Acta2. Third, AAV-mediated restoration of SAMP1 in the lung attenuates the same endpoints. Fourth, at the cellular level, SAMP1-deficient AT2 cells show reduced 53BP1 together with increased PAI-1 and p16^INK4a^, linking the loss of this inner nuclear membrane protein to an impaired DNA damage response and a senescent, profibrotic state.

Our findings place SAMP1 within a growing framework in which the nuclear envelope i regarded not merely as a structural boundary but as an active participant in genome maintenance. Chromatin anchored at the inner nuclear membrane exhibits distinct repair kinetics, damaged loci relocate to nuclear pore complexes, and LINC complex components couple the nucleus to cytoskeletal motors to mobilize broken chromosomes [16,17,46]. Correspondingly, perturbation of inner nuclear membrane proteins delays double-strand break repair and produces premature ageing phenotypes [18,46]. SAMP1 is well positioned to participate in this axis: it is targeted to the inner nuclear membrane through CxxC zinc-finger motifs, binds lamin A/C, emerin and the LINC components SUN1 and SUN2, and contributes to peripheral heterochromatin anchoring [19,20,21,50]. A reduction of 53BP1 in SAMP1-deficient AT2 cells is therefore consistent with a model in which loss of SAMP1 compromises the capacity of the epithelium to mount an effective damage response. We note importantly, however, that 53BP1 was measured under baseline, unchallenged conditions and that γH2AX was unchanged; our data therefore establish an alteration in damage-response machinery, but do not by themselves demonstrate accumulation of unrepaired DNA breaks. Experiments in which DNA damage is induced experimentally and its resolution followed over time — for example by quantifying γH2AX and 53BP1 foci clearance after BLM, etoposide or irradiation — will be required to distinguish defective repair from altered steady-state protein expression.

The concomitant induction of p16^INK4a^ and PAI-1 in SAMP1-deficient AT2 cells is significant because AT2 cell senescence is now recognized as an active driver, rather than a passive consequence, of lung fibrosis. Induction of p53-dependent senescence selectively in adult AT2 cells produces spontaneous progressive fibrosis that is reversed when senescence is blocked or senescent cells are cleared [8,42], and telomere dysfunction confined to AT2 cells is likewise sufficient to cause lung remodeling and fibrosis [11,43]. Senescent AT2 cells elaborate a SASP that acts in an autocrine and paracrine manner to promote myofibroblast differentiation and matrix deposition [10,12,13,42]. PAI-1 occupies a distinctive position within this programme: it is inducible by TGF-β, and it functions not only as a marker but as a mediator of senescence, being required for TGF-β1-induced AT2 cell senescence and SASP secretion, while its deletion or pharmacological inhibition attenuates BLM-induced fibrosis [12,44,45]. The presence of elevated PAI-1 in SAMP1-deficient AT2 cells therefore suggests a plausible route by which epithelial SAMP1 loss translates into a profibrotic tissue environment.

An unexpected and informative aspect of our results is the cellular selectivity of the SAMP1 change. Whereas SAMP1 was clearly reduced in AT2 cells from fibrotic lungs, its level in primary lung fibroblasts was comparable between saline- and BLM-treated mice. This asymmetry aligns with the current view that the epithelium, rather than the mesenchymal or immune compartment, initiates fibrogenesis [5,8,40], and it argues that the pro-fibrotic consequence of SAMP1 loss is best understood as an epithelial-intrinsic defect. It also has practical implications: the fibroblast data make it unlikely that the exacerbated fibrosis observed in Samp1^AT2-KO^ mice is secondary to a mesenchymal effect of the conditional allele.

The therapeutic implication of our work is that SAMP1 is not only a susceptibility factor but a tractable target. AAV-mediated delivery has emerged as a credible platform for pulmonary gene transfer, with several serotypes efficiently transducing the airway and alveolar epithelium in preclinical models [30,47]. That a single intratracheal administration of AAV-Samp1 reduced fibrosis scores and lowered Tnfα and Acta2 expression in our hands provides proof of concept that augmenting SAMP1 is beneficial in this model. A senescence-directed reading of these data would suggest that preserving SAMP1 function, or intervening downstream at the level of PAI-1 or senescent cell clearance, merits further evaluation [42,44,47].

## Conclusions

SAMP1/TMEM201 is selectively reduced in alveolar type 2 cells during bleomycin-induced pulmonary fibrosis; its deletion in AT2 cells aggravates fibrosis, whereas its restoration ameliorates it. At the cellular level, SAMP1 deficiency is accompanied by reduced 53BP1 and by induction of the senescence-associated and profibrotic markers p16^INK4a^ and PAI-1. These results position SAMP1 as a guardian of AT2 cell integrity against DNA damage–induced senescence and identify the SAMP1–AT2 senescence axis as a candidate contributor to pulmonary fibrogenesis and a potential target for epithelial-directed therapy.

## Declarations

### Preprint status

This manuscript has been deposited as a preprint and has not been peer-reviewed. The findings have not been confirmed by independent review and should not be used to guide clinical practice or health-related behaviour. A version of record, if published, will be linked from the preprint record.

### Author contributions

L.A.S.: Conceptualization, Methodology, Investigation, Formal analysis, Writing – original draft. D.A.B.: Methodology, Investigation, Formal analysis, Visualization, Writing – review and editing. Both authors reviewed and approved the final manuscript and agree to be accountable for all aspects of the work.

### Data and materials availability

All data supporting the conclusions are presented in the main figures and in Supplementary Tables 1 and 2. Additional materials and reagents are available from the corresponding author upon reasonable request, subject to institutional and national regulations.

### Use of generative AI and AI-assisted tools

Generative AI and large language model tools were used during manuscript preparation for language editing, reference formatting, figure-legend standardisation and assembly of the manuscript and declaration sections. No AI tool was used to generate, fabricate or analyse primary research data, and no AI tool is listed as an author. All scientific content, experimental values, citations and interpretations were checked against the original laboratory records and source publications, and were reviewed, edited and approved by the authors, who take full responsibility for the accuracy, integrity and originality of the work.

## Supplementary Tables

*SAMP1/TMEM201 protects alveolar type 2 cells from DNA damage-associated senescence and restrains pulmonary fibrosis*

**Supplementary Table 1.**
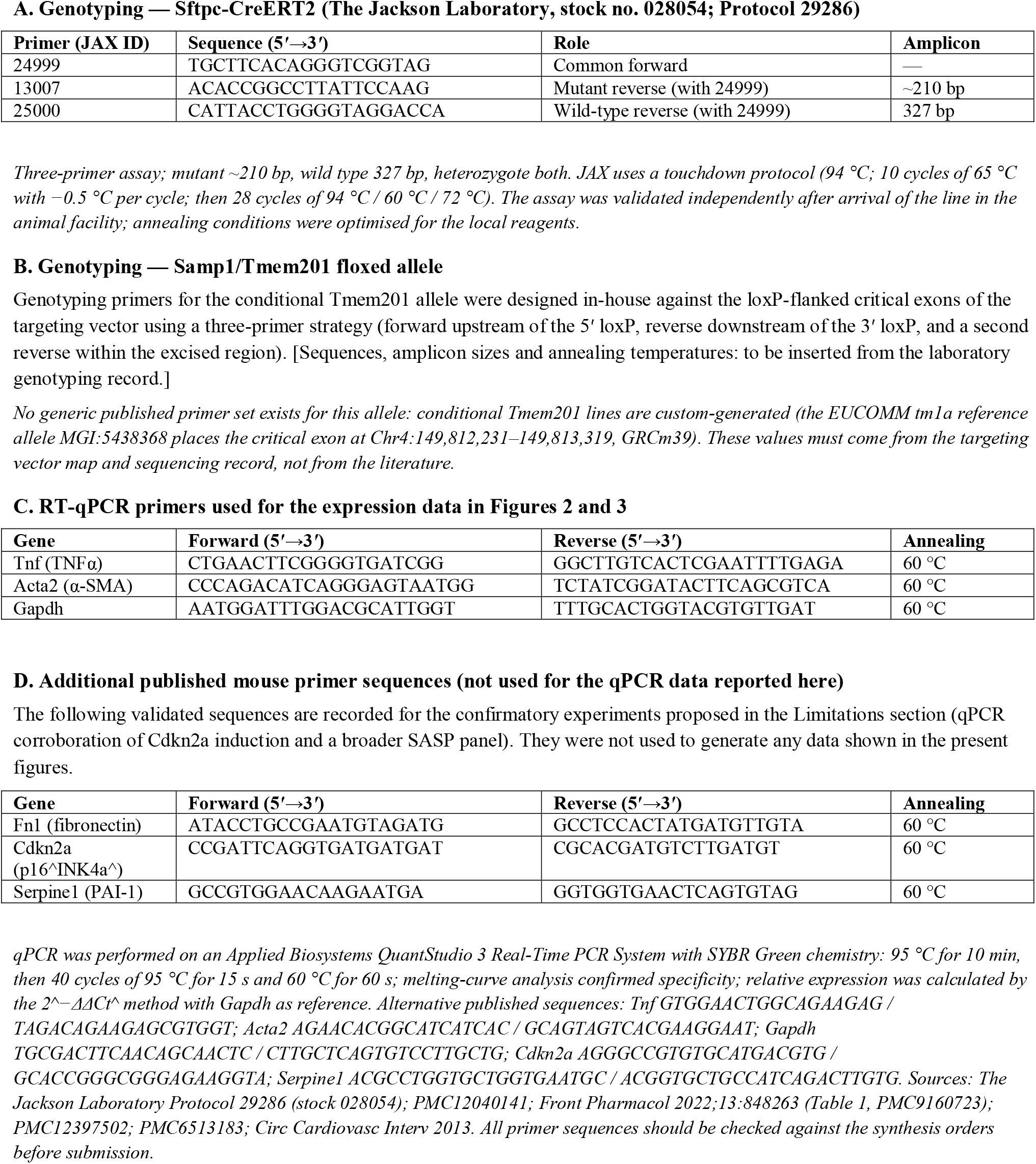
Primer sequences.

**Supplementary Table 2.** Antibodies and working dilutions.

| **Target** | **Supplier / Cat. no.** | **RRID** | **Host / clonality** | **WB dilution** | **Use** |
|---|---|---|---|---|---|
| SAMP1 / TMEM201 | Proteintech 87333-1-RR | AB_3745473 | Rabbit, recombinant mAb | 1:10,000 (1:5,000–1:50,000) | WB; IF co-staining |
| Fibronectin | Abcam ab2413 | AB_2262874 | Rabbit, polyclonal | 1 µg/mL | WB |
| α-SMA (ACTA2) | Sigma-Aldrich A2547 (1A4) | AB_476701 | Mouse, monoclonal IgG2a | 1:1,000–1:2,000 | WB |
| 53BP1 | Novus Biologicals NB100-304 | AB_10003037 | Rabbit, polyclonal | 1–2 µg/mL | WB |
| γH2AX (Ser139) | Cell Signaling 9718 (20E3) | AB_2118009 | Rabbit, monoclonal | 1:1,000 | WB |
| PAI-1 (SERPINE1) | Proteintech 66261-1-Ig (1H4A5) | AB_2881648 | Mouse, monoclonal IgG1 | 1:5,000–1:50,000 | WB |
| p16^INK4a^ | Abcam ab108349 (EPR1473) | AB_10858268 | Rabbit, monoclonal | 1:4,000 | WB |
| pro-SP-C | Merck Millipore AB3786 | AB_91588 | Rabbit, polyclonal serum | 1:500 | WB; IF |
| GAPDH | Cell Signaling 5174 (D16H11) |  | Rabbit, monoclonal | 1:1,000 | WB loading control |
| HRP goat anti-rabbit IgG | Cell Signaling 7074 |  | Goat, polyclonal | 1:1,000–1:3,000 | WB secondary |
| HRP goat anti-mouse IgG | Cell Signaling 7076 |  | Goat, polyclonal | 1:1,000–1:3,000 | WB secondary |

